# Morphological heterogeneity of human astrocytes in cerebral organoids

**DOI:** 10.1101/2025.09.18.675430

**Authors:** Kaitlin Szederkenyi, Aaron Au, Liliana Attisano, Martin Oheim, Christopher M. Yip

## Abstract

Astrocytes are a type of glial cell in the central nervous system responsible for modulating synaptic transmissions, tissue repair, maintaining homeostasis, and are therefore implicated in many neurological diseases. Human cortical astrocytes are more structurally complex, larger, and have unique subtypes in comparison to the commonly studied rodent cortical astrocytes. As access to human cortical tissue is sparse, cerebral organoids (COs) derived from human pluripotent stem cells have emerged as a promising in vitro model for studying the human cortex. Astrocyte subtypes unique to humans are recapitulated in COs but have not been quantitatively assessed (1, 2). In this study, we characterized human astrocytes in situ in sliced COs cultured at the air-liquid interface (ALI-COs). By 4 months of age, ALI-COs express many mature astrocyte markers and showed increasing levels of glial fibrillary acidic protein (GFAP) with longer culture durations. By employing immunostaining, tissue clearing, morphological reconstruction, and unsupervised clustering analysis, three major GFAP+ astrocyte subtypes were identified in ALI-COs. All subtypes exhibited greater morphological complexity than their mouse counterparts, as revealed by increased branching and longer branch extensions. However, consistent with the mid-gestation fetal stage of the ALI-COs, astrocytes did not fully recapitulate the complexity observed in adult human astrocytes, which are known to continue maturation postnatally.

## Introduction

The macro- and micro-anatomical differences between the human and rodent brain are well established (3, 4). However, due to restricted access to healthy human brain tissue, rodents remain the predominant model for neuroscience research. While studies of the rodent cortex have yielded valuable insights into glial function, fundamental differences between rodent and human cortical architecture limit the direct translational applicability of many findings (3-6).

Historically, cortical astrocytes were considered a homogenous population. However, it has recently been shown that there is both morphological and transcriptomic heterogeneity across the cortex, suggesting varying functional roles (7-9). In the human cortex, four morphologically distinct glial fibrillary acidic protein-positive (GFAP+) astrocyte subtypes have been characterized: interlaminar astrocytes in layers 1–2, protoplasmic in layers 2–6, varicose in layers 5–6, and fibrous astrocytes in the white matter (10).

Interlaminar astrocytes (ILA) in the human cortex have cell bodies located in the pial or subpial, regions, and they extend long processes down to layers 3–4. Comparative analyses across 46 mammalian species, from rodents to primates, indicate that human ILAs exhibit the highest structural complexity (11). In rodents, pial astrocytes are rudimentary and lack long processes. Human protoplasmic astrocytes are also markedly distinct from their rodent counterparts, being 2.6-fold larger, possessing 10-fold more primary branches, and propagating calcium waves four times faster in response to ATP and glutamate stimulation (10).

Varicose astrocytes, characterized by 1-5 GFAP+ fibers up to 1 mm in length with regularly spaced varicosities, have been reported exclusively in hominid brains. However, their presence exhibits inter-individual variability, as they were not identified in all human subjects examined, raising questions about whether they represent a distinct astrocyte subtype or a disease phenotype (10, 12). Human fibrous astrocytes are also larger than those in rodents, with more extensive processes that display significant interdigitation with neighbouring fibrous astrocytes (10). As morphological differences between human and rodent astrocytes continue to be elucidated, there is a growing need for physiologically relevant live models to move beyond static morphological descriptions and explore the functional properties of distinct human astrocytes.

Human, including primary and stem-cell-derived, astrocytes are commonly studied using two-dimensional (2D) culture systems. However, astrocytes cultured in 2D often fail to recapitulate the complex morphology observed in three-dimensional (3D) environments, they frequently adopt a reactive phenotype and lack critical signals from other cells in the surrounding tissue. Live resected human tissue provides a more physiologically relevant alternative. However, in the rare cases where available for research, it is often comprised of fragments resected from patients undergoing surgery for a pathological condition such as epilepsy or tumours. This raises concerns regarding relevance to healthy physiology and makes it difficult to obtain samples that span cortical layers and of specific ages.

Cerebral organoids (CO) derived from human pluripotent stem cells have emerged as a powerful model for studying neurodevelopment and disease. The transcriptomic changes associated with CO aging have been characterized by multiple groups, and their functionality has been assessed through electrophysiology and neuronal calcium imaging (1, 13-16). Astrocytes dissociated from COs and cultured in 2D display key astrocytic functions, including glutamate uptake, synaptosome phagocytosis, and promoting synaptic formation (17). Moreover, interlaminar, protoplasmic, and fibrous astrocyte subtypes have been identified in a slice CO model and in glia-enriched COs transplanted into mouse brain, but the observed morphologies have not been characterized (1, 2).

In this study, we characterized the morphologies of GFAP+ cells identified in 5-month-old air-liquid interface COs (ALI-COs) into defined major human subtypes and compared these morphologies to adult mouse and human astrocytes. We demonstrate that all subtypes exhibit greater morphological complexity than their mouse counterparts, as shown by increased branching and longer branch extension. However, ALI-CO astrocytes did not fully recapitulate the complexity observed in adult human astrocytes, likely reflecting their mid-gestational developmental stage, as human astrocytes are known to continue maturation postnatally and possibly into adolescence (18, 19). Our findings highlight the potential of ALI-COs as a valuable complementary model to enhance the translation relevance of rodent-based astrocyte studies in human health and disease.

## Results

### Characterization of the GFAP+ cell population in ALI-COs

In a previous study, transcriptomic analysis of COs derived from the same H9 embryonic stem cell line and cultured using the same protocol as in this study revealed the emergence of a cluster with mature astrocytic identity at 18 weeks, which continues to expand with extended culture duration (16). However, at this age, fewer than 1% of cells express GFAP, and approximately 2% express the calcium binding protein S100B, a consensual astrocyte marker. By 24 weeks, the expression of these markers increased to 5% and 10%, respectively (16). This prompts the question of the optimal age at which to study astrocytes in COs, to achieve a sufficient number of mature astrocytes, while minimizing the culture duration. Prolonged culture leads to the development of necrotic cores within the COs due to restricted oxygen and nutrient diffusion to the innermost regions. Previous studies have shown that slicing COs and continuing culture at the air-liquid interface (ALI) improves both health and maturation (13, 20). Accordingly, we adopted an ALI-CO culture method (Figure 1). COs were matured for 90 days in free-floating spinning culture, as described by (16), then sectioned and maintained under ALI conditions for extended culture durations.

**Figure 1.**
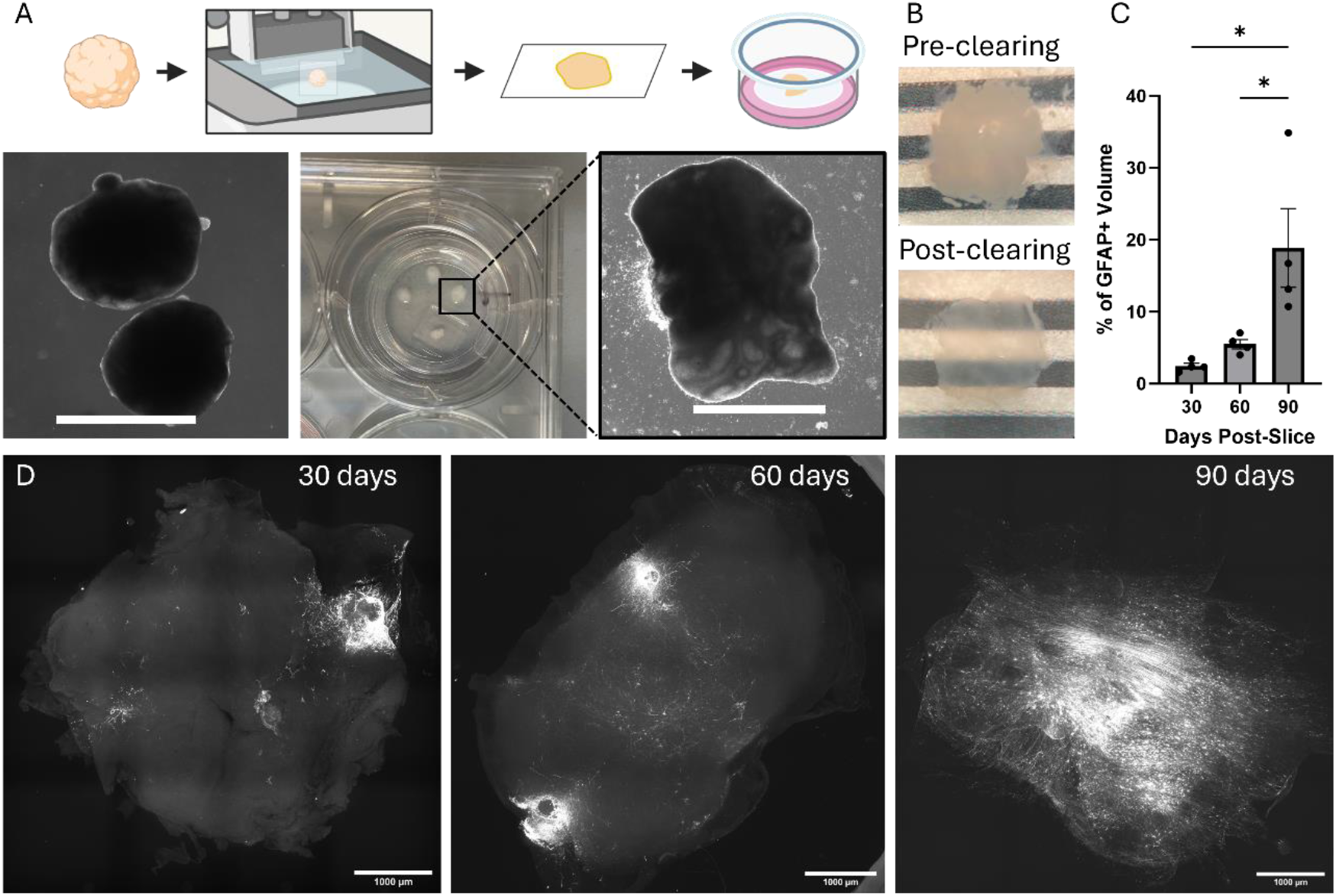
GFAP+ cells occupy more territory over time in ALI-COs. A) *Top*: Cerebral organoids (COs) are sliced at 90 days-old and culture is continued on air-liquid interface (ALI) membrane inserts (created with BioRender.com). *Bottom:* Images show corresponding steps, from *left* to *right*, COs in spin culture, COs sliced on an ALI membrane insert, and zoomed in on an ALI-CO. Scale bars are 2 mm. B) ALI-CO from first image of D) before and after optical clearing with Ubiclear. C) % of ALI-CO volume occupied by GFAP+ cells and their processes (N = 4, error bars = SEM, one-way ANOVA followed by Tukey’s test, * = p < 0.05). D) Representative ALI-COs immunolabelled for GFAP after 90 days in spin culture followed by 30, 60, and 90 days, respectively, in ALI culture. Images are MIPs of 300 μm acquired on a Nikon CREST spinning disk confocal microscope.

At 4-months old (90 days in spin + 30 days in ALI culture), ALI-COs co-expressed the astrocyte markers Vimentin and PAX6, as well as specifically mature astrocyte markers S100B, AQP4, ALDH1L1, and Connexin 43, along with GFAP (Supplementary Figure 1). Qualitatively, GFAP+ populations did not have any apparent co-localization with either the mitotic marker KI67 or the radial glia marker BLBP, suggesting that the majority of GFAP+ cells are not radial glia at this timepoint. As anticipated, GFAP+ cells interact structurally with the neuronal population as revealed by TUBB3 staining, but do not co-express the neuronal marker (Supplementary Figure 1).

GFAP+ cell maturity can also be assessed spatially, based on whether cells are still heavily bundled at ventricular zones, moving outwards, or mostly located past these zones. Intact ALI-COs were optically cleared, and full volumes were imaged using spinning disk confocal microscopy (Figure 1). After 90 days in spin + 30 days in ALI culture, most GFAP+ cells were still heavily bundled near ventricular-like zones with few single cells dispersed throughout the tissue. GFAP+ cell mass accounted for less than 2.5% of the ALI-CO volume. After 60 days, there was an increase in GFAP+ single cells dispersed throughout the tissue displaying astrocytic morphology, and GFAP+ cell mass increased to 5.5%. By 90 days, there was a larger increase in volume occupied by GFAP+ cells and their processes now occupied an average of 18.9% of the ALI-COs volume (Figure 1). At this age, astrocyte processes became densely packed, thereby complicating the characterization of the morphology of individual cells using immunolabelling. Consequently, for subsequent analysis, we used 5-month-old (90 days in spin + 60 days in ALI culture) ALI -COs to quantify the morphological traits of ALI-CO astrocytes.

### ALI-COs display morphologically distinct GFAP+ cell subtypes

GFAP+ cells in 5-month-old fixed, cleared, and immunolabelled ALI-COs were manually classified as protoplasmic, fibrous, ILA, or progenitor-like based on established morphological criteria (9, 10, 21). Varicose astrocytes were not observed in the ALI-COs. ILAs were identified by the presence of a soma located within the superficial layer at the tissue edge in the XY-plane, with one or more process extending inward toward deeper regions. Protoplasmic astrocytes were identified as having a bushy structure and many primary branches of similar length. Fibrous astrocytes were identified by having fewer primary branches and extending their process far from their cell body. GFAP can also be expressed across populations of radial glia and intermediate progenitor cells populations, both of which are known to be present in ALI-COs at this developmental stage. Therefore, GFAP+ cells exhibiting rudimentary morphology, characterized by 1–3 single processes extending from their soma, were classified broadly as progenitor-like cells (Figure 2). Morphological parameters of each group were quantified as defined in Supplementary Table S1. Of the four identified groups, protoplasmic astrocytes, exhibited the largest number of primary branches, followed by fibrous and ILA, and then progenitor-like cells with the least branches (proto = 22.27 ± SD 8.77, fibro = 12.43 ± 5.18, ILA = 9.56 ± 4.82, PC = 2 ± 1.07). To assess branching complexity, several parameters were measured, including the maximum Sholl intersections at a single radius, total number of paths, Horton Stahler number, and maximum branch fractal dimension. Protoplasmic astrocytes displayed the highest degree of branching across these metrics, except for a non-significant difference in mean branch fractal dimension compared to fibrous (max Sholl intersections: proto = 34.43 ± 16.17, fibro = 17.21 ± 7.31, ILA = 13.31 ± 6.76, PC = 2.75 ± 1.83, # of paths: proto = 70.4 ± 39.09, fibro = 42.82 ± 28.14, ILA = 22.38 ± 9.67, PC = 3.88 ± 2.7; Horton Stahler number: proto = 4.03 ± 0.42, fibro = 3.68 ± 0.61, ILA = 3.06 ± 0.44, PC = 1.75 ± 0.71; branch fractal dimension: proto = 1.3 ± 0.13, fibro = 1.25 ± 0.11, ILA = 1.15 ± 0.06, PC = 1.08 ± 0.05). Fibrous astrocytes and ILAs exhibited similar branching patterns, although fibrous astrocytes demonstrated a modestly greater complexity, particularly in number of paths and branch fractal dimension. In terms of spatial footprint, fibrous, ILAs, and PCs displayed more elongated morphologies than protoplasmic astrocytes, as reflected by larger maximum Sholl radii, and an increased longest shortest path (max Sholl radius (μm): proto = 70.93 ± 22.7, fibro = 127.4 ± 52.07, ILA = 118.5 ± 74.55, PC = 123 ± 27.69; longest shortest path (μm): proto = 105.1 ± 33.47, fibro = 163.6 ± 61.83, ILA = 141 ± 84.23, PC = 154.9 ± 33.03). When comparing cable length (i.e., the sum of all branch lengths), a parameter that incorporates both branching complexity and elongation, protoplasmic and fibrous astrocytes were not significantly different but they were both larger than ILA, and all three were significantly larger than PCs (cable length (in μm): proto = 1615 ± 893.9, fibro = 1217 ± 738.2, ILA = 636.6 ± 221.2, PC = 227.1 ± 77.1). Protoplasmic astrocytes exhibited the highest degree of roundness, and were more symmetrical than fibrous astrocytes as seen by the skewness of their branching (roundness: proto = 0.75 ± 0.1, fibro = 0.56 ± 0.13, ILA = 0.59 ± 0.15, PC = 0.35 ± 0.08; skewness: proto = 0.48 ± 0.24, fibro = 0.86 ± 0.52, ILA = 1.02 ± 0.86, PC = 0.39 ± 0.24). Overall, fibrous and ILA astrocytes showed similar morphometric profiles, differing significantly only in the number of paths and Horton-Strahler number (Figure 2). The primary distinguishing feature between these two subtypes was spatial: interlaminar astrocytes were consistently localized to the superficial edges of the ALI-COs. Notably, this morphological similarity between fibrous and interlaminar astrocytes has also been reported in human cortical tissue (22).

**Figure 2.**
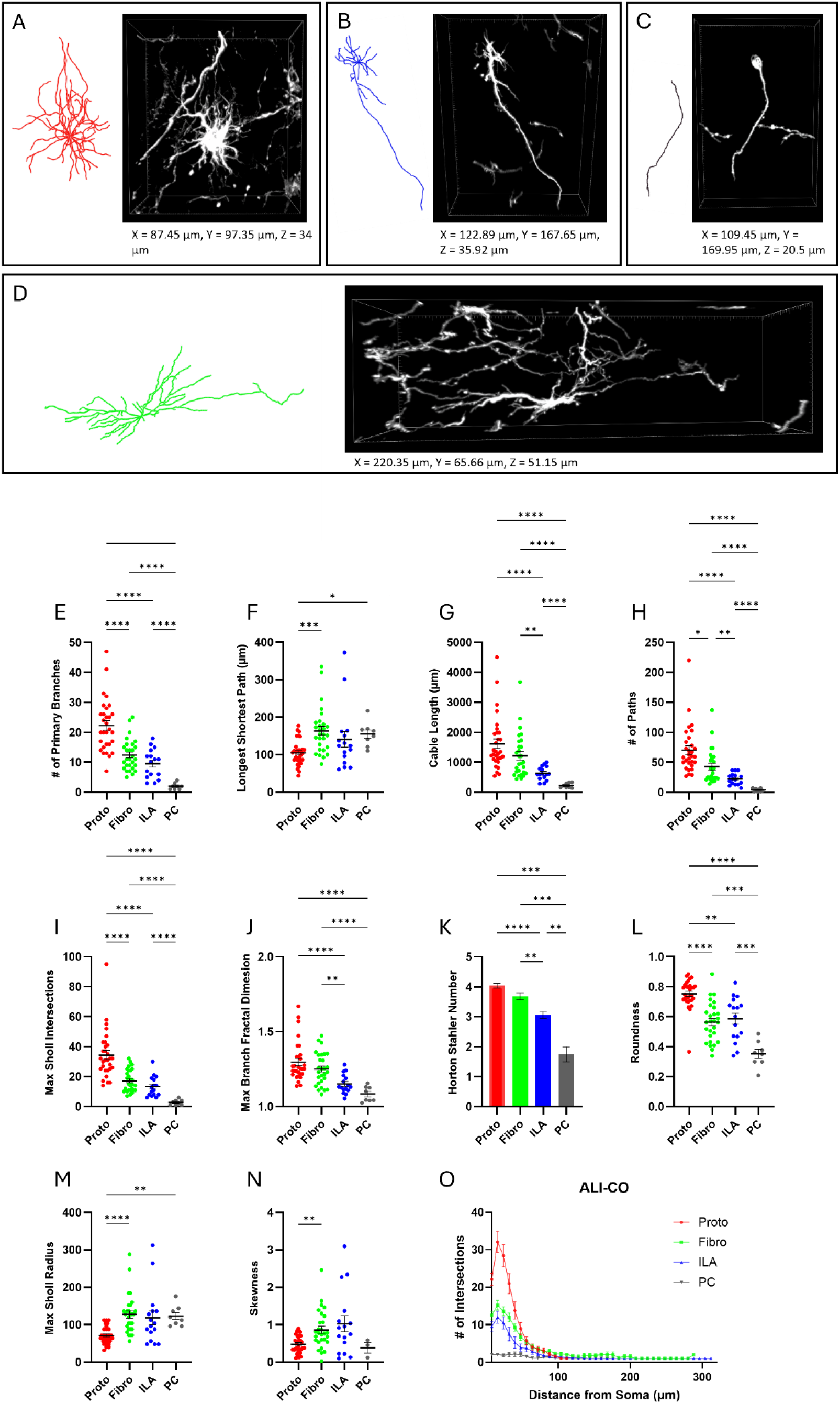
GFAP+ astrocyte subtypes in ALI-COs have distinct morphological features. Representative Gtree reconstruction and corresponding 3D maximum projections of A) protoplasmic, B) interlaminar, and D) fibrous astrocytes; and C) a progenitor-like cell identified in 5-month-old ALI-COs. Dimensions correspond to the 3D ROI box sizes. Protoplasmic (proto), fibrous (fibro), interlaminar (ILA), and progenitor-like cells (PC) were characterized based on the E) number of primary branches, F) longest shortest path, G) cable length, H) total # of paths, I) maximum number of intersections of a single Sholl radius, J) maximum branch fractal dimension, K) Horton Stahler number, L) roundness, M) maximum Sholl radius distance from soma, N) skewness of branching distribution, and O) Sholl profile of entire reconstructions. Proto n = 30, fibro n = 28, ILA n = 16, PC n = 8, N = 6 ALI-COs (n = # of cells, N = # of tissue samples). Black bars indicate the mean and error bars are SEM. * = p < 0.05, ** = p < 0.01, *** = p < 0.001, **** = p < 0.0001, Brown-Forsythe and Welch’s ANOVA, followed by Dunnett’s T3 test.

To validate our manual classification of the GFAP+ cell morphological subtypes, the cells were classified using a Gaussian Mixed Model (GMM) unsupervised clustering algorithm, based on the parameters quantified in Figure 2 (23). With four defined classes, the GMM achieved an overall classification accuracy of 94%. PCs and ILA were distinguished with the highest accuracy, both yielding F1 scores of 100%. Greater overlap was observed between protoplasmic and fibrous astrocytes, nevertheless these subtypes were classified with 92% and 90% accuracies, respectively (Figure 3). While fibrous and ILA morphologies exhibit similarities, incorporating the superficial localization of ILAs as a clustering parameter enabled their separation into distinct classes with high accuracy. Although GMM assigns class membership probabilistically, all cells were classified with high confidence, as seen in Figure 3B.

**Figure 3.**
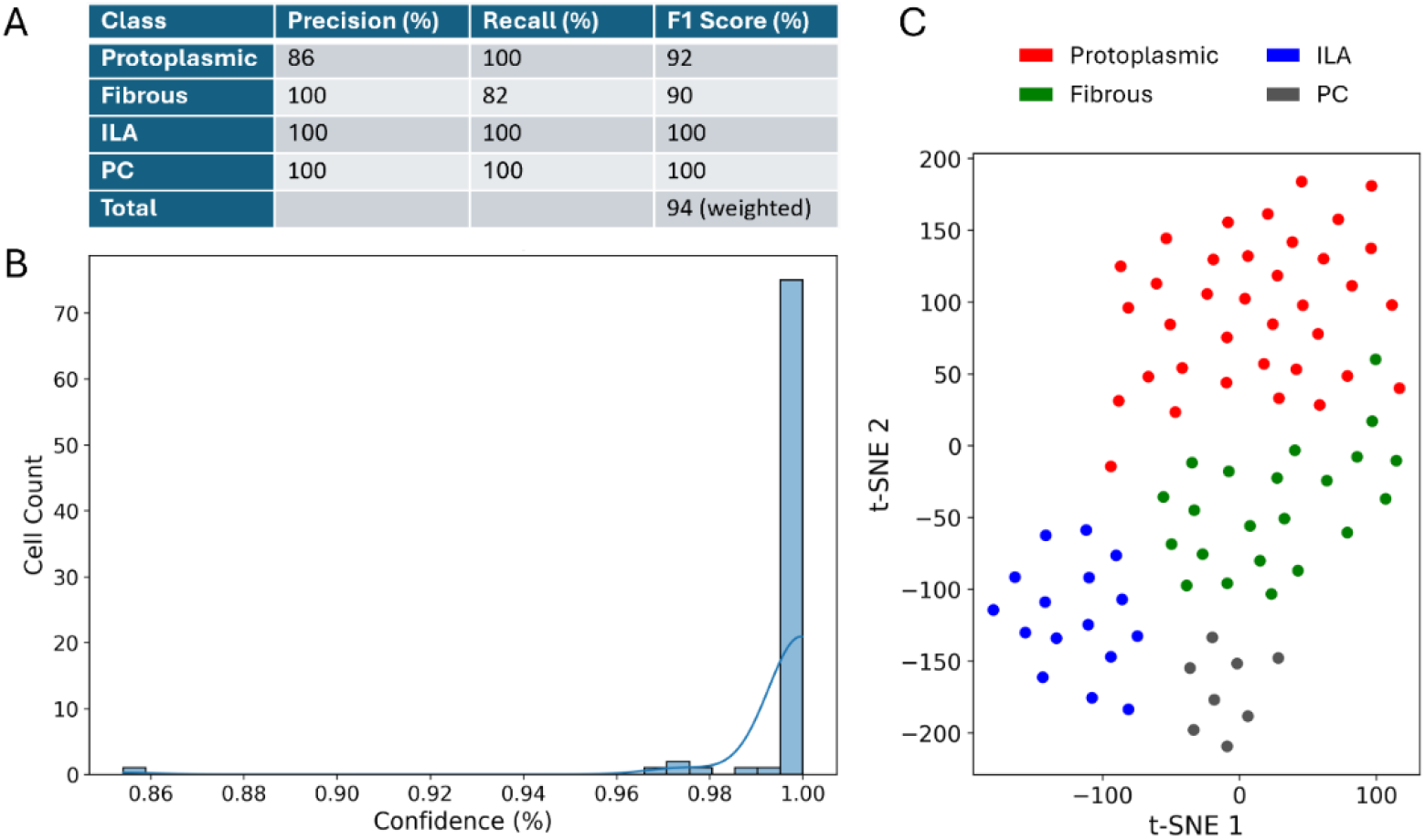
GFAP+ subtypes in ALI-COs are classified with high accuracy by unsupervised Gaussian mixed model clustering. A) Evaluation of model accuracy, B) percentage confidence of a cell belonging to the cluster in which it was classified, and C) a t-SNE plot illustrating the classification result. n = 82 cells, N = 6 ALI-COs.

### ALI-CO astrocytes are more complex than those found in adult mice

For comparison of ALI-CO astrocytes to those in the mouse cortex, postnatal day 30 (P30) mouse cortical slices were cleared and immunolabeled for GFAP. Individual GFAP+ astrocytes were then reconstructed and quantified using the same methods applied to the ALI-CO cells. For consistency, all ROIs were obtained from the somatosensory cortex. Pial astrocytes were identified as GFAP+ cells with their soma in the pial layer, protoplasmic astrocytes as GFAP+ cells located between layers 2–6, and fibrous astrocytes as those located in the cortical white matter.

We found that across all morphological subtypes, ALI-CO astrocytes had more primary branching than their mouse counterparts (mouse # of primary branches: proto = 4.82 ± 1.25, fibro = 5.67 ± 2.57, pial = 3.6 ± 1.34) (Figure 4). ALI-CO astrocytes also displayed longer cable lengths and longest shortest paths, implying that they are larger in size and elongate their branches further from the soma (mouse cable length (μm): proto = 584 ± 124.6, fibro = 500.1 ± 170.2, pial = 175.6 ± 44.5; longest shortest path (μm): proto = 60.84 ± 8.29, fibro = 60.84 ± 8.29, pial = 45.06 ± 24.61). Interestingly, only protoplasmic and ILA had more higher-order branching than their mouse counterparts, whereas fibrous astrocytes had similar levels of higher order branching (mouse max Sholl intersections: proto = 19.09 ± 3.48, fibro = 15.92 ± 4.42, pial = 7.6). ALI-CO fibrous astrocytes did, however, have more total paths, as a result of more primary branches (mouse # of paths: proto = 29.82 ± 4.92, fibro = 24.83 ± 9.45, pial = 11 ± 1.41). Mouse astrocytes were consistently more symmetrical than those of ALI-COs, and fibrous and pial astrocytes were rounder (mouse skewness: proto = 0.2 ± 0.14, fibro = 0.19 ± 0.09, pial = 0.27 ± 0.16; roundness: proto = 0.78 ± 0.06, fibro = 0.8 ± 0.05, pial = 0.78 ± 0.08). Overall, ALI-CO astrocytes are larger, extend more primary branches, and are more asymmetrical than their mouse counterparts (Figure 4).

**Figure 4.**
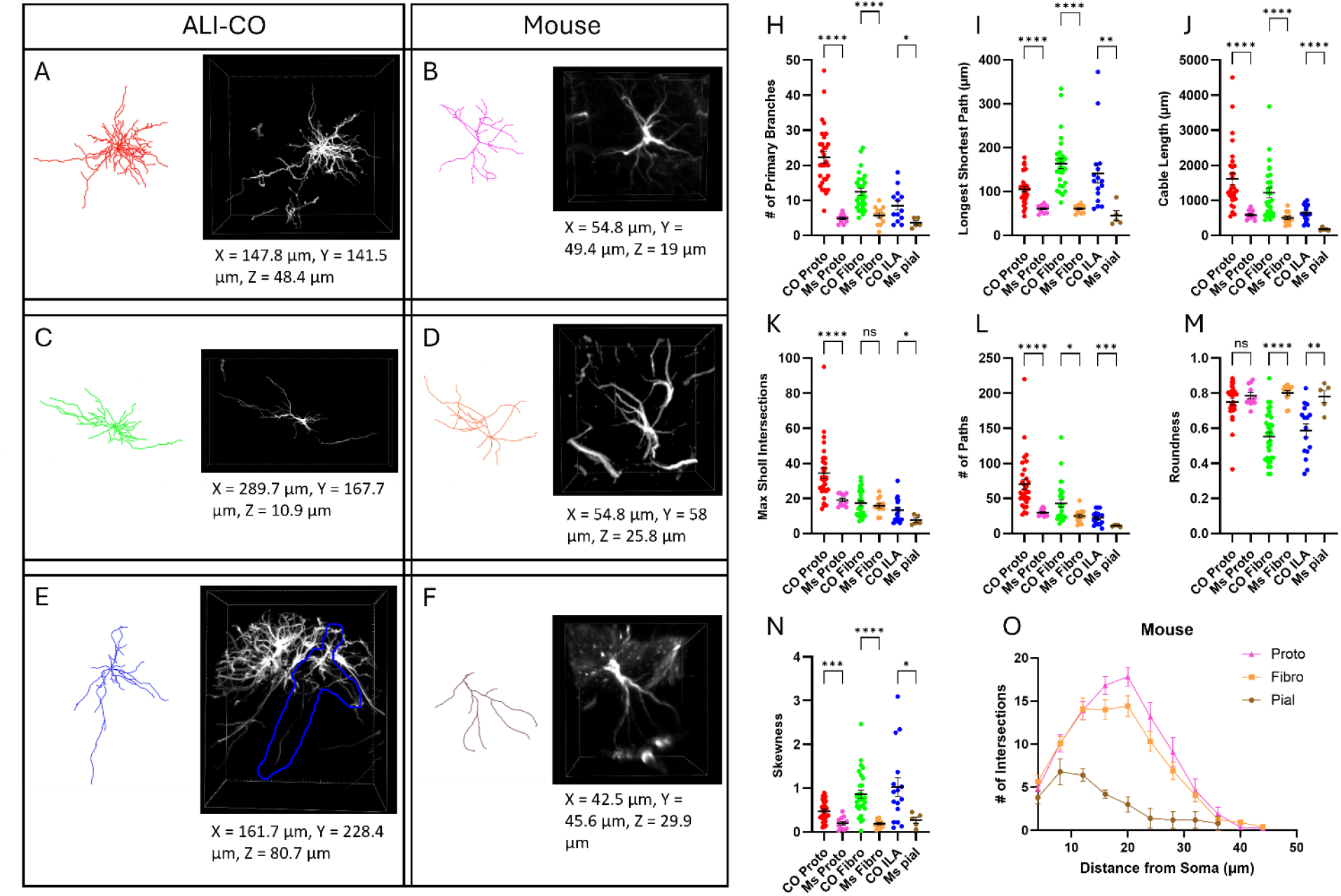
ALI-COs astrocytes are larger and more branched than their mouse counterparts. Representative Gtree reconstructions and corresponding 3D ROIs of A) ALI-CO and B) P30 mouse protoplasmic; C) and D) fibrous; and E) and F) ILAs. Dimensions correspond to the 3D ROI box sizes. Protoplasmic (proto), fibrous (fibro), interlaminar (ILA) astrocytes belonging to ALI-COs (CO) or Mice (Ms) were characterized based on H) the number of primary branches, I) longest shortest path, J) cable length, K) maximum number of intersections of a single Sholl radius, L) total # of paths, M) roundness, N) skewness of branching distribution, and O) Sholl profile of entire reconstructions. CO proto n = 30, CO fibro n = 28, CO ILA n = 16, N = 6 ALI-COs. Ms proto n = 11, Ms fibro n = 12, Ms pial n = 5, N = 3 cortical slices from 3 animals. Statistics are as described in Figure 2.

### GFAP+ protoplasmic astrocytes of ALI-COs extend their branches further but have less branching than human adult protoplasmic astrocytes

Postmortem human cortical tissue obtained from a single 47-year-old female with no known neurological conditions was cleared and immunostained for GFAP. Densely packed ILAs were identified in layer 1 with processes extending into layer 4. Protoplasmic astrocytes were found in layers 3–6, with large end feet at blood vessels, while fibrous astrocytes formed a dense mesh of filaments in the white matter (Figure 5). Varicose astrocytes were not identified; however, the data was collected only from one subject, and inter-subject variability has been reported (12). ILA and fibrous astrocytes were too dense to distinguish single cells sufficiently for reconstruction, however, protoplasmic astrocytes were reconstructed and quantified alongside their ALI-CO and mouse counterparts.

**Figure 5.**
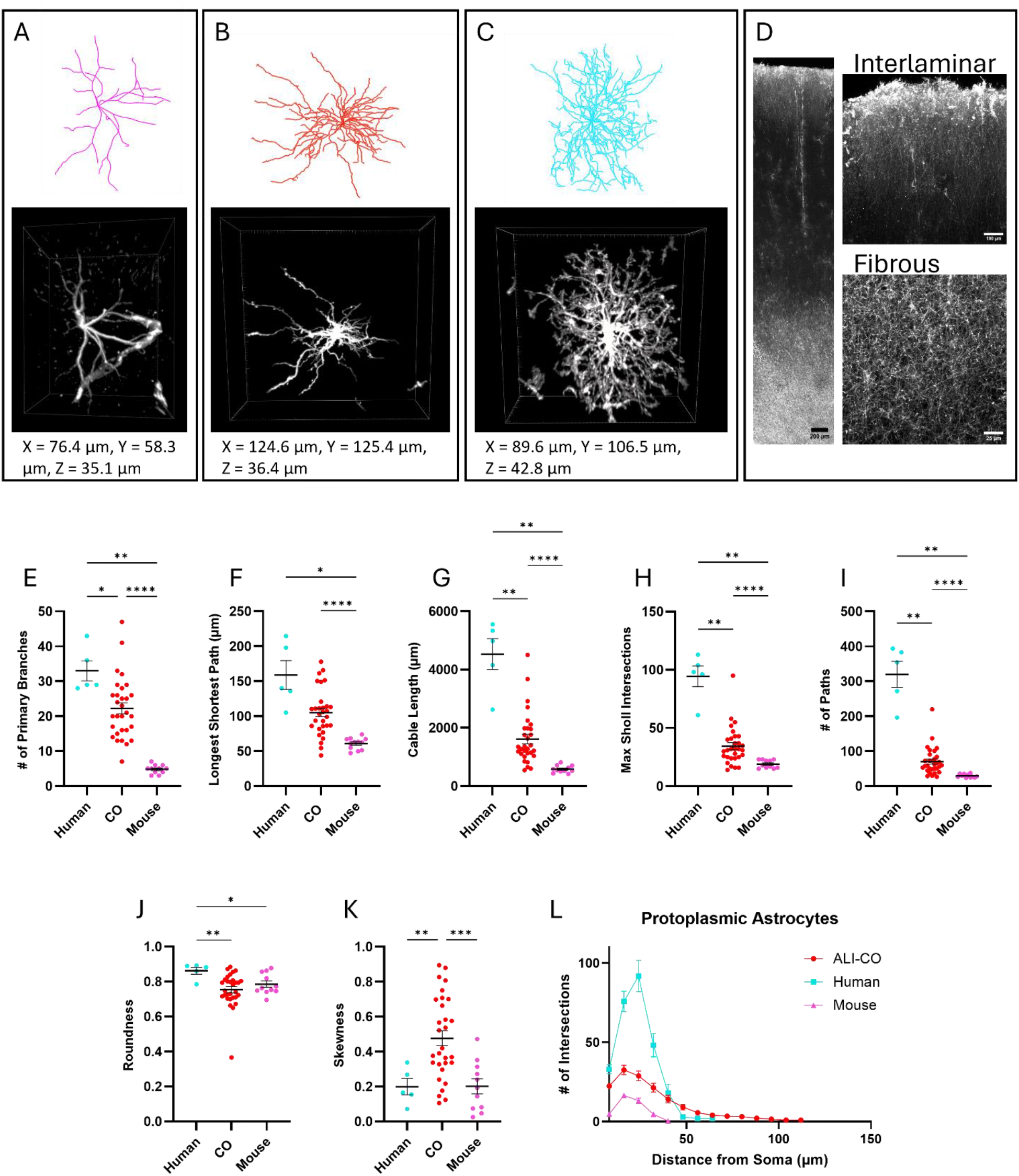
GFAP+ ALI-CO protoplasmic astrocytes are less branched but extend their branches further than those of adult human astrocytes. Representative Gtree reconstructions and corresponding 3D ROIs of GFAP+ cells in A) P30 mouse, B) ALI-CO, and C) adult human protoplasmic astrocytes. Dimensions correspond to the 3D ROI box sizes. D) GFAP labelling across human cortical layers, followed by interlaminar (layer 1-2) and fibrous (white matter) regions, presented as MIPs of 100 μm. Protoplasmic astrocytes of adult human, ALI-CO, and P30 mouse were characterized based on the E) number of primary branches, F) longest shortest path, G) cable length, H) maximum number of intersections of a single Sholl radius, I) total # of paths, J) roundness, K) skewness of branching distribution, and L) Sholl profile of entire reconstructions. Human n = 5 cells, N = 3 cortical slices from 1 individual; CO n = 30, N = 6 ALI-COs; mouse n = 11, N = 3 cortical slices from 3 animals. Statistics are as described in Figure 2.

The human adult protoplasmic astrocytes had more primary branches than ALI-COs (human = 33 ± 6.44), higher branching complexity as indicated by a larger number of paths (human = 319.8 ± 84.8), max Sholl intersections (human = 94.4 ± 19.81), and cable length (human = 4528 ± 1190 μm). They also elongated their branches further, demonstrated by a greater longest shortest path (human = 158.7 ± 45.95 μm). However, the maximum Sholl radius was not significantly larger than that of mouse protoplasmic astrocytes, and significantly smaller than that of ALI-COs. This may reflect the presence of less linear, more tortuous processes. Thus, taken together, protoplasmic ALI-CO astrocytes exhibit greater branching complexity than their mouse counterparts, though not to the extent observed in adult human astrocytes. Notably, 5-month-old ALI-CO astrocytes achieve elongations lengths comparable to those of human astrocytes. It is possible, however, that ALI-COs are lacking systemic in vivo cues that normally constrain branch extension in healthy individuals.

## Discussion

While transcriptomic profiling of COs generated from various protocols has been performed across developmental timepoints and compared to human and mouse cortical transcriptomes, in situ morphological characterization of CO astrocytes remains limited (1, 2, 17, 24). In this study, ALI-COs were found to contain 3 GFAP+ morphologically distinct astrocyte subtypes, in addition to GFAP+ progenitor-like cells having a more rudimentary morphology. While subtypes are distinct enough to be accurately classified by an unsupervised clustering algorithm, considerable intra-subtype variability was observed across morphological metrics. It is likely that with larger sample sizes and the inclusion of additional astrocyte markers, further heterogeneity within these groups would be revealed. Integrative approaches that capture both morphological and transcriptomic profiles, such as Patch-seq (25), hold great potential for deepening our understanding of the astrocyte diversity within COs, but are beyond the scope of this work.

Our findings reveal that ALI-CO astrocytes are more complex than those in the mouse cortex but they remain still less elaborate than their adult human counterparts, as they have less primary and higher order branching than human astrocytes. Notably, protoplasmic astrocytes in the ALI-Cos extended their branches further than those in both the mouse and human cortex. This extended branching may reflect a lack of in vivo cues that limit branch extension, or possibly a compensatory response to the absence of vasculature in the organoid environment, since in vivo protoplasmic astrocytes interact extensively with blood vessels to facilitate transport and maintain homeostasis. Additionally, while varicose astrocytes were not found in our ALI-COs, this phenotype was recently identified in glia-enriched COs transplanted into the mouse cortex (2). The observation of varicose astrocytes following transplantation is particularly interesting, as this subtype has not yet been observed in COs maintained entirely in vitro. This raises the possibility that specific environmental cues present in the in vivo brain microenvironment, but absent in organoid cultures, may be required for the emergence or maturation of this astrocytic phenotype.

Although the sample size was limited, the observed morphologies of fibrous and ILA in the human cortex, as well as the quantified morphology of protoplasmic astrocytes, were broadly consistent with previous reports. The average number of primary branches in protoplasmic astrocytes (33 primary branches) closely matched values reported by (10) (37.5 primary branches) and the measured elongation was within the previously described range of 100–400 μm (10). However, it remains important to note that this data was derived from a single individual, and the exact sampling location within the prefrontal cortex was undefined. Additionally, the tissue was fixed postmortem, and the time between death and fixation is unknown. Despite these limitations, the reasonable agreement between our measurements and prior studies supports both the validity of the methodology used and suitability of the obtained human cortical tissue as benchmark for comparative analysis with ALI-CO astrocytes.

It is also important to emphasize that the human cortical tissue analyzed in this study is adult, while COs are representative of the developing brain. While an ideal comparison would involve astrocytes in mid- to late gestation human fetal cortical tissue in situ, such samples were not available for this study and, to our knowledge, detailed morphological analyses in this context have not yet been reported. Furthermore, our study provides a snapshot of ALI-CO astrocytes at 5 months of age, however, astrocyte maturation is a dynamic process that continues with extended culture duration, as indicated by a marked increase in GFAP+ astrocyte density by 6 months of age. Nevertheless, the degree of morphological complexity, specifically the increased branching and size, of ALI-COs astrocytes at 5 months is already striking when compared to P30 mouse cortical astrocytes. Also, the higher astrocyte density at greater ages makes the identification and morphometry of individual cells more difficult, if not impossible. Sparse cell labeling via viral transduction instead of immunolabeling could, however, allow for the morphological characterization at these stages.

While the many functions of fibrous and protoplasmic astrocytes have been identified in rodents and 2D culture, the function of ILAs remains unknown. Due to the drastic difference between primate interlaminar and mouse pial astrocytes, it has been hypothesized that these cells may be involved in the enhanced cognitive abilities of primates (11). In humans, ILAs are reported to be more numerous, longer (extending up to 1 mm), and overall more complex than those observed in New World monkeys, with greater inter-individual variability as well (26, 27). In this context, ALI-COs offer a promising platform to further study these primate-specific cells and elucidate the function of this enigmatic astrocyte sub-population.

In summary, our study provides a detailed in situ morphological characterization of major astrocytes subtypes in ALI-COs, and comparison with mouse and human cortical astrocytes in situ. This comprehensive characterization of astrocyte morphology provides a critical baseline for probing more specific questions about astrocyte heterogeneity, enables functional studies of identified astrocyte populations, and informs the identification and refinement of strategies for therapeutic manipulation in a human-specific 3D model.

## Materials and Methods

### CO and ALI-CO culture

COs were cultured as described in (16). At 3 months of age, COs were placed in cryomolds (Fisher Scientific NC9511236) and excess media removed. The COs were then embedded in 4% low melting point agarose (ThermoFisher Scientific 16520050). A Leica VT 1200S vibratome set to run at 0.45 m/s with a forward displacement of 1 mm/s was used to prepare 300-µm thick slices of the embedded COs. End slices were discarded, and the middle sections were placed on Millicell cell culture inserts (Millipore Sigma PICM0RG50) for ALI-CO culture (Figure 1A). During their culture, 800 µl cerebral organoid differentiation medium (composition described in (16)) was placed under the inserts, and the medium was changed every 72 h. ALI-COs were incubated at 37°C and 5% CO_2_.

### Mouse cortex

All experiments followed European Union (EU) and institutional (CNRS) guidelines for the care and use of laboratory animals (Council Directive 2010/63/EU and CEUA approval number 001-01-2023, respectively). This study used a total of 3 adult (postnatal day 30) C57BL/6J mice. Fixed cortices were obtained from Dr. Antonin Singer at SPPIN, who performed the fixation as described in (28). A vibratome (Leica VT 1000S) was used to prepare 300 µm-thick sagittal slices.

### Human cortex

Postmortem sections of prefrontal cortex were fixed in 4% paraformaldehyde (PFA) for 48 h then put into phosphate buffered saline (PBS). The donor was a 47-year-old female with no known neurological conditions. The tissue was obtained through collaboration with Dr. Eleonora Aronica, MD, PhD, Department of Neuropathology, Meibergdreef 9, 1105 AZ, Amsterdam, The Netherlands. Cortex sections were then equally sliced on a vibratome to 300 µm thickness.

### Cryosections

COs and ALI-COs were rinsed with PBS without Ca^2+^ and Mg^2+^ (PBS -/-), fixed in 4% PFA overnight at 4°C, then washed 3 × 30 min with PBS -/-. They were soaked in 15% sucrose solution for 2 h, then 30% sucrose overnight at 4 °C. They were then transferred to a cryomold containing optimal cutting temperature (OCT) compound (VWR 95057-838) and flash-frozen in liquid nitrogen. Samples were sectioned to 14-µm thickness using a cryostat (Leica CM3050S), mounted on Superfrost plus microscope slides (ThermoFisher Scientific 22-037-246) (3–6 slices per slide) and heated for 15 min at 45°C. Slices were stored at -20°C.

### Cryosection immunofluorescence microscopy

Slices were warmed to room temperature, then circles were drawn around slices of interest with a Readyprobes hydrophobic barrier pap pen (ThermoFisher Scientific R3777). Slices were rehydrated in PBS -/- for 5 min, permeabilized with 0.5% Triton X-100 (Millipore Sigma 9002-93-1) in PBS-/- for 30 min, then rinsed 3X with PBS-/-. Slices were blocked with 5% donkey serum (Millipore Sigma D9663) in PBS-/- for 30 min. Primary antibodies were diluted in antibody solution (PBS-/- with 1% donkey serum) at the following concentrations: anti-GFAP (Thermo fisher scientfic 14-9892-80) 1:500, anti-TUBB3 (ThermoFisher Scientific MA1-118X) 1:200, anti-KI67 (Proteintech 27309-1-AP) 1:100, anti-PAX6 (Proteintech 12323-1-AP) 1:250, anti-OLIG2 (Proteintech 13999-1-AP) 1:100, anti-Vimentin (Proteintech 10366-1-AP) 1:250, anti-Aquaporin 4 (16473-1-AP) 1:200, anti-Myelin basic protein (MBP) (Proteintech 10458-1-AP) 1:50, anti-Connexin 43 (Millipore Sigma C6219) 1:200, anti-ALDH1L1 (ThermoFisher Scientific PA5-78822) 1:300, anti-S100B (Abcam ab52642) 1:400, anti-BLBP (Abcam ab32423) 1:50. Slices were incubated with primary antibodies overnight in sealed box containing wet paper towel to prevent evaporation at 4°C. The next day, slices were washed with PBS-/- 3 × 10 min, then incubated with secondary antibodies in antibody solution at the following concentrations: Alexa fluor 555 goat anti-mouse (ThermoFisher Scientific A21424) 1:500, Alexa Fluor 488 chicken anti-rabbit (ThermoFisher Scientific A21441) 1:500, and Alexa Fluor 647 donkey anti-rabbit (ThermoFisher Scientific A31573) 1:500; and 4′,6-diamidino-2-phenylindole (DAPI) (60 µl/ml) for 1 h at room temperature. Slices were washed 3 × 10 min, then 1 drop (30 µl)/slice of Prolong gold antifade mountant (ThermoFisher Scientific P10144) was applied and the slices were covered with a coverslip.

Imaging was done on either a laser-scanning confocal microscope (Nikon Eclipse Ti2 AX) using a Plan-Apochromat 20X 0.75 NA objective, with excitation at 405 nm, 488 nm, 561 nm, or 640 nm laser lines at 3–35% laser power and a gain of 10–30; or a laser-scanning confocal Zeiss LSM710 using a Plan-Apochromat 20X 0.8 NA or Plan-Apochromat 63X 1.4 NA oil objective, and excited at 405 nm, 488 nm, 561 nm at 5–30% laser power. Full volumes were acquired at Nyquist sampling conditions and are presented as maximum intensity projections (MIPs).

### Tissue clearing and full volume immunofluorescence microscopy

Mouse and human cortex samples were depigmented in Ubiclear (Idylle-labs) depigmentation solution for 1 h at 37°C with constant agitation. All subsequent steps were performed under the same conditions. Tissues were placed in 2 ml microcentrifuge tubes, rinsed with 1.5 ml of tris(hydroxymethyl)aminomethane (tris) buffer (0.2 M, pH 8), then dehydrated in 1.5 ml of the following methanol (MeOH) dilutions for 15 min each (ALI-CO and mouse) or 30 min (human): 50% MeOH/tris, 80% MeOH/tris, 100% MeOH, 2 × 80% MeOH/dimethyl sulfoxide (DMSO) (Millipore Sigma 67-68-5), 80% MeOH/tris, 50% MeOH/tris, then for mouse and ALI-CO only 2 × tris, followed by 2 × 1% Triton X-100/tris. Human samples required an additional protein denaturing step with the following acetone dilutions for 30 min each: 50% acetone/tris, 2 × 100% acetone, 2 × 50% acetone/tris, 2 × tris, 1% Triton X-100/tris. Samples were then soaked in 1 ml penetration buffer (0.2% Triton X-100, 0.3 M glycine, and 20% DMSO in tris) for 1 h and then blocking buffer (0.2% Triton X-100, 6% donkey serum, and 10% DMSO in tris) for 5 h (ALI-CO and mouse) or overnight (human). Primary antibodies at concentrations previously defined were applied in antibody buffer (3% donkey serum and 5% DMSO in wash buffer (2% Tween20 (Thomas scientific C987C37), 0.1 mg/ml heparin (Millipore Sigma 9041-08-1), in tris)) overnight. Samples were washed 5 × 30 min with washing buffer, and secondary antibodies were applied overnight at previously defined concentrations in antibody buffer. Samples were again washed 5 × 30 min with washing buffer, in tris buffer for 30 min, then optically cleared in Ubiclear clearing agent (Idylle-labs) for 30 min (mouse), 1 h (CO), or 3 h (human), and finally into storage/imaging solution (60% 2,2’-thiodiethanol (Millipore Sigma 111-48-8) and 3% triethylenediamine (Millipore Sigma 280-57-9), in tris).

Samples were imaged either on a laser-scanning confocal microscope (Nikon Eclipse Ti2 AX) using a 20X 0.95 NA water immersion objective, excited with 405 nm, 488 nm, 561 nm, and 640 nm laser lines at 5–30% laser power; or a laser-scanning confocal Zeiss LSM710 using a Plan-Apochromat 20X 0.8 NA or Plan-Apochromat 63X 1.4 NA oil objective using the 405 nm, 488 nm, 561 nm, and 633 nm laser lines at 5–30% laser power; or a spinning disk confocal Nikon eclipse Ti2-E crest X equipped with a Prime 95B camera (Teledyne Photometrics) using a Plan-Apochromat 10X 0.45 NA or 40X Apochromat long working distance 1.15 NA water immersion objective, excited at 405 nm, 473 nm, 545 nm, and 635 nm at 10–40% laser power, and exposure time of 50–150 ms.

### Image analysis

3D ROIs of single cells were extracted from the fixed, cleared, and immunostained full ALI-CO volumes. ROIs were preprocessed by applying a custom python script which segmented the fluorescence signal from the background. First, the raw images were background subtracted using a gaussian filter (σ = 25 pixels). A mask was then produced by thresholding the background subtracted image to values above the 3rd standard deviation in intensity. Small objects were removed from the mask, and then the objects were dilated by 1 pixel. The mask was applied to the raw images which were then converted to 8-bit images. Images were then loaded into GTree, an automated reconstruction algorithm (29), and single-cell morphologies were reconstructed. However, Gtree occasionally failed to trace small branches with dim intensity or erroneously reconstructed branches that were not present. Therefore, to ensure accuracy, all reconstructions were subsequently reviewed and manually corrected using the Simple Neurite Tracer (SNT) Fiji plugin (30). Raw unprocessed images were used to identify regions requiring correction. Morphological parameters were then measured using SNT.

### Gaussian mixed model (GMM) clustering

Clustering was performed by applying GMM from the Python SciKit-learn library (23) using the following morphological parameters: # of primary branches, roundness, Sholl skewness, # of paths, max Sholl intersections, max Sholl radius, proximity to edge, cable length, max branch fractal dimension, and the Horton Stahler number. All parameter distributions were first scaled using PowerTransformer to stabilize variance. GMM was run with the following settings: number of user defined clusters = 4, covariance type = full, and random state = 42. Results were visualized using t-SNE, also from the scikit-learn python toolbox (23). Data was scaled using PowerTransformer and t-SNE was run with the following settings: perplexity = 30, learning rate = 500, number of iterations = 1000, random state = 42. The accuracy of the model was measured by its F1 score:

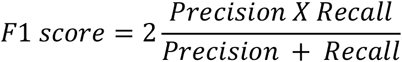

Where:

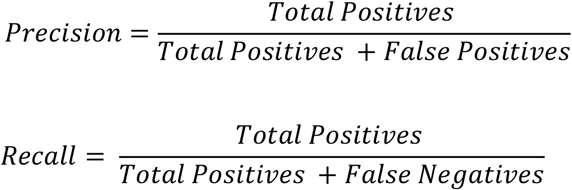

### Statistics

Statistics were conducted through Graphpad Prism 10.4.1. Normality was tested for parameter distributions through D’Agostino and Pearson, Anderson-Darling, and Shapiro-Wilk tests, as well as visual inspection of QQ plots. For normally distributed data, one-way ANOVA followed by Tukey’s post-hoc test, or Brown-Forsythe and Welch ANOVA followed by Dunnett’s T3 test were applied (if there was unequal variance across groups). Supplementary Table 2 provides full statistical reports for all data sets.

## Acknowledgments

We thank Hui Zhang and Dulce Carrillo Valenzuela at the Applied Organoid Core at the University of Toronto for their help with generation and maintenance of organoids. We also thank Dr. Antonin Singer and Dr. Eleonora Aronica for tissue preparations, and Brigitte Delhomme for assistance with tissue clearing. This work was supported by Natural Sciences and Engineering Research Council of Canada [RGPIN-2015-043 to C.M.Y] and a CNRS-UofT Twin Fellowship to M.O. and C.Y. This Project was also made possible with the financial support of Health Canada, through the Canada Brain Research Fund, an innovative partnership between the Government of Canada (through Health Canada) and Brain Canada, and of the Krembil Foundation [to L.A].

## Abbreviations

ALI-CO: Air liquid interface cerebral organoid
CO: Cerebral organoid
ALI: Air liquid interface
DMSO: Dimethyl sulfoxide
Fibro: Fibrous
GFAP: Glial fibrillary acidic protein
GMM: Gaussian Mixed Model
ILA: Interlaminar astrocyte
MIP: Maximum intensity projection
PBS: Phosphate buffered saline
PC: Progenitor cell
PFA: Paraformaldehyde
Proto: Protoplasmic

## Author Contributions

K.S., M.O., C.M.Y., and L.A. designed research; K.S. performed the research, K.S. and A.A. analyzed data; K.S., M.O., and C.M.Y. wrote the paper.

