## Supplementary for "Morphological heterogeneity of human astrocytes in cerebral organoids"

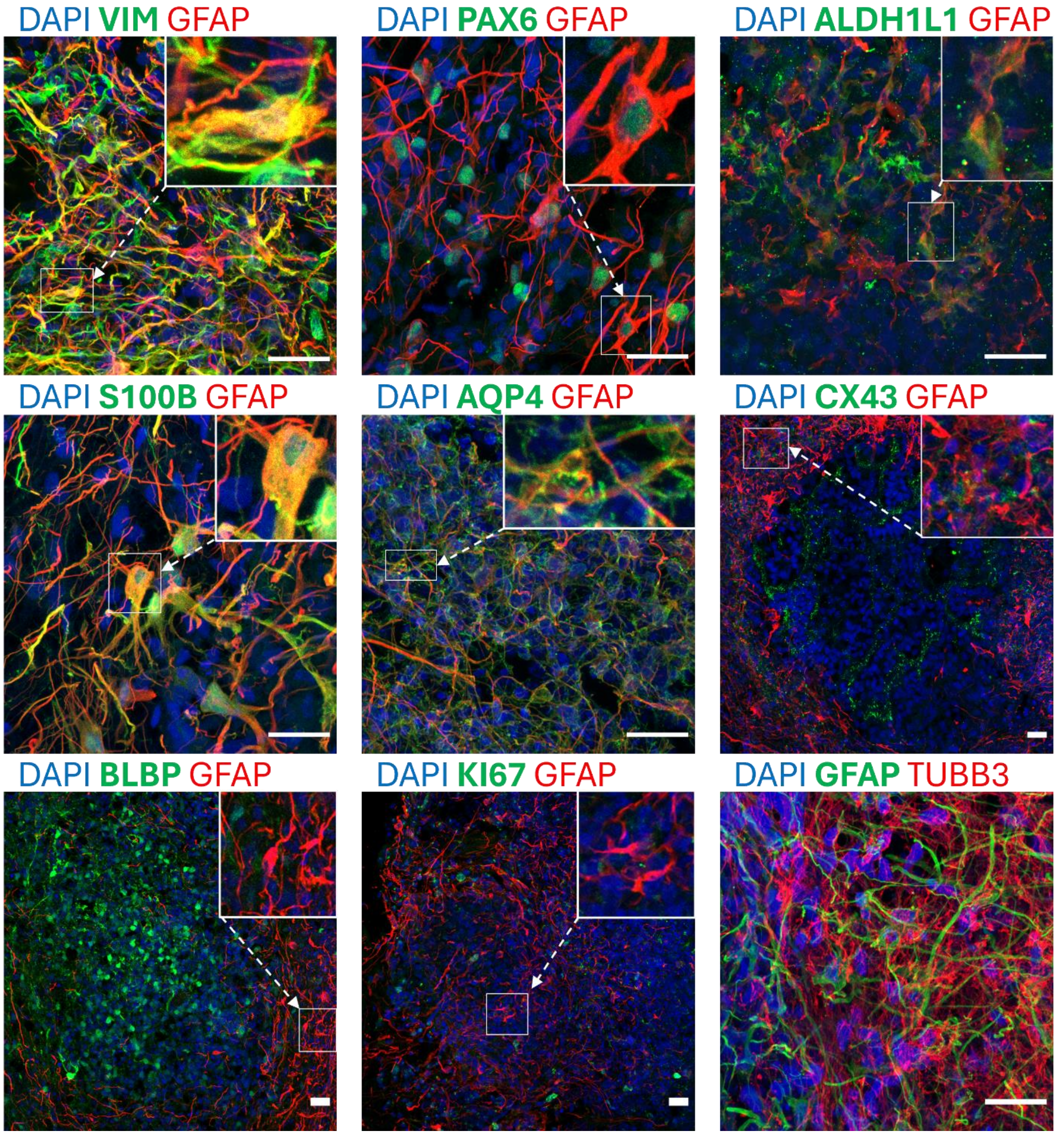


**Supplementary Figure 1. GFAP+ cell populations co-localize with astrocyte markers but not proliferation markers.** MIPs of ALI-COs (90 days in spin culture + 30 days in ALI culture) cryosectioned to 14 µm, and immunostained for GFAP and astrocyte markers (Vimentin, PAX6, ALDH1L1, S100B, AQP4, and CX43), proliferation markers (BLBP and KI67), and neuronal marker TUBB3. Boxes on top right are enlarged ROIs. All scale bars = 25 µm.

**Supplementary Table 1. Description of morphological measurements calculated through SNT (30).**

| | |
|---|---|
| Roundness | How closely the shape of a 3D convex hull resembles a sphere (4π Area / Perimeter$^2$) |
| # of primary branches | # of branches attached to the soma, calculated from the first Sholl radius. |
| Max Sholl radius | Maximum distance branches extend from soma (rounded to the nearest Sholl radius) |
| Max Sholl intersections | Maximum number of branches intersecting a single Sholl radius. |
| Sholl skewness | Symmetry of the branching distribution relative to the soma, the larger the value the more asymmetrical. |
| Longest shortest path (µm) | Measure of how "spread out" the cell is. Longest distance along an arbour from one branch tip to another (longest geodesic). (µm) |
| Horton-Stahler root number | Measure of branching complexity in a tree-like structure. Starts with 1 at the tips and increases as more branches of the same value merge towards the soma. |
| Branch fractal dimension | Describes the complexity of branching structure. The slope of the log-log plot of path distance vs. Euclidean distance. |
| Cable length (µm) | Sum of all path lengths (µm) |
| # of paths | Total number of paths |
| Proximity to edge | Whether the cell soma is located in the outmost XY layer of the CO. |

**Supplementary Table 2. Detailed statistical report for all figures.**

| Figure and Panel | Statistical test | Degrees of freedom (DFn) | Degrees of freedom (DFd) | F value or W value | Actual P value |
|---|---|---|---|---|---|
| Figure 1C | Brown-Forsythe test | 2 | 9 | 2.009 | 0.19 |

| | | | | | |
|---|---|---|---|---|---|
| | Ordinary one-way ANOVA | 2 | 9 | 7.481 | 0.0122 |
|  | **Tukey's multiple comparisons test**<br><br>**Test** **Adjusted P value** **q** **DF**<br>30 vs. 60 0.7831 0.9552 9<br>30 vs. 90 0.0136 5.142 9<br>60 vs. 90 0.0384 4.187 9 |  |  |  |  |
| Figure 2E | Brown-Forsythe ANOVA | 3 | 63.45 | 41.44 | <0.0001 |
| | Welch's ANOVA | 3 | 39.89 | 80.63 | <0.0001 |
|  | **Dunnett's T3 multiple comparisons test**<br><br>**Test** **Adjusted P value** **t** **DF**<br>Proto vs. Fibro <0.0001 5.242 47.60<br>Proto vs. ILA <0.0001 6.342 43.92<br>Proto vs. PC <0.0001 12.32 31.91<br>Fibro vs. ILA 0.3549 1.847 33.31<br>Fibro vs. PC <0.0001 9.936 32.83<br>ILA vs. PC <0.0001 5.993 17.73 |  |  |  |  |
| Figure 2F | Brown-Forsythe ANOVA | 3 | 38.39 | 5.355 | 0.0035 |
| | Welch's ANOVA | 3 | 26.36 | 9.081 | 0.0003 |
|  | **Dunnett's T3 multiple comparisons test**<br><br>**Test** **Adjusted P value** **t** **DF**<br>Proto vs. Fibro 0.0004 4.436 40.94<br>Proto vs. ILA 0.5020 1.634 17.57<br>Proto vs. PC 0.0170 3.774 11.16<br>Fibro vs. ILA 0.9161 0.9414 24.37<br>Fibro vs. PC 0.9948 0.5298 22.25<br>ILA vs. PC 0.9917 0.5780 21.32 |  |  |  |  |
| Figure 2G | Brown-Forsythe ANOVA | 3 | 60.35 | 19.38 | <0.0001 |
| | Welch's ANOVA | 3 | 41.63 | 47.03 | <0.0001 |
|  | **Dunnett's T3 multiple comparisons test**<br><br>**Test** **Adjusted P value** **t** **DF**<br>Proto vs. Fibro 0.3402 1.856 55.20 |  |  |  |  |

| | | | | | |
|---|---|---|---|---|---|
| | Proto vs. ILA | <0.0001 | 5.678 | 35.15 | |
| | Proto vs. PC | <0.0001 | 8.388 | 30.54 | |
| | Fibro vs. ILA | 0.0027 | 3.865 | 34.62 | |
| | Fibro vs. PC | <0.0001 | 6.961 | 28.94 | |
| | ILA vs. PC | <0.0001 | 6.641 | 20.57 | |
| Figure 2H | Brown-Forsythe ANOVA | 3 | 57.71 | 26.21 | <0.0001 |
| | Welch's ANOVA | 3 | 40.56 | 55.93 | <0.0001 |
| | **Dunnett's T3 multiple comparisons test** | | | | |
| | **Test** | **Adjusted P value** | **t** | **DF** | |
| | Proto vs. Fibro | 0.0184 | 3.099 | 52.69 | |
| | Proto vs. ILA | <0.0001 | 6.373 | 35.14 | |
| | Proto vs. PC | <0.0001 | 9.239 | 30.00 | |
| | Fibro vs. ILA | 0.0073 | 3.500 | 36.50 | |
| | Fibro vs. PC | <0.0001 | 7.209 | 28.65 | |
| | ILA vs. PC | <0.0001 | 7.121 | 19.04 | |
| Figure 2I | Brown-Forsythe ANOVA | 3 | 53.43 | 39.10 | <0.0001 |
| | Welch's ANOVA | 3 | 40.19 | 63.86 | <0.0001 |
| | **Dunnett's T3 multiple comparisons test** | | | | |
| | **Test** | **Adjusted P value** | **t** | **DF** | |
| | Proto vs. Fibro | <0.0001 | 5.283 | 40.98 | |
| | Proto vs. ILA | <0.0001 | 6.209 | 42.33 | |
| | Proto vs. PC | <0.0001 | 10.48 | 31.56 | |
| | Fibro vs. ILA | 0.3906 | 1.788 | 33.45 | |
| | Fibro vs. PC | <0.0001 | 9.480 | 33.86 | |
| | ILA vs. PC | <0.0001 | 5.836 | 18.86 | |
| Figure 2J | Brown-Forsythe ANOVA | 3 | 75.13 | 17.17 | <0.0001 |
| | Welch's ANOVA | 3 | 33.34 | 21.64 | <0.0001 |
| | **Dunnett's T3 multiple comparisons test** | | | | |
| | **Test** | **Adjusted P value** | **t** | **DF** | |
| | Proto vs. Fibro | 0.6056 | 1.468 | 55.18 | |
| | Proto vs. ILA | <0.0001 | 5.068 | 43.72 | |
| | Proto vs. PC | <0.0001 | 7.111 | 30.72 | |
| | Fibro vs. ILA | 0.0024 | 3.841 | 41.95 | |
| | Fibro vs. PC | <0.0001 | 6.127 | 25.76 | |

| | | | | | |
|---|---|---|---|---|---|
| | ILA vs. PC 0.0651 2.818 17.25 | | | | |
| Figure 2K | Brown-Forsythe ANOVA | 3 | 27.82 | 38.37 | <0.0001 |
| | Welch's ANOVA | 3 | 24.96 | 35.29 | <0.0001 |
| | **Dunnett's T3 multiple comparisons test**<br><br>**Test** **Adjusted P value** **t** **DF**<br>Proto vs. Fibro 0.0766 2.568 47.00<br>Proto vs. ILA <0.0001 7.247 28.99<br>Proto vs. PC 0.0001 8.743 8.320<br>Fibro vs. ILA 0.0026 3.850 39.49<br>Fibro vs. PC 0.0002 7.002 10.19<br>ILA vs. PC 0.0040 4.801 9.834 | | | | |
| Figure 2L | Brown-Forsythe ANOVA | 3 | 52.36 | 29.27 | <0.0001 |
| | Welch's ANOVA | 3 | 27.28 | 44.12 | <0.0001 |
| | **Dunnett's T3 multiple comparisons test**<br><br>**Test** **Adjusted P value** **t** **DF**<br>Proto vs. Fibro <0.0001 6.237 49.96<br>Proto vs. ILA 0.0031 4.058 22.20<br>Proto vs. PC <0.0001 11.34 12.21<br>Fibro vs. ILA 0.9952 0.5247 28.22<br>Fibro vs. PC 0.0003 5.404 17.12<br>ILA vs. PC 0.0004 4.913 21.21 | | | | |
| Figure 2M | Brown-Forsythe ANOVA | 3 | 34.75 | 7.475 | 0.0005 |
| | Welch's ANOVA | 3 | 24.75 | 15.26 | <0.0001 |
| | **Dunnett's T3 multiple comparisons test**<br><br>**Test** **Adjusted P value** **t** **DF**<br>Proto vs. Fibro <0.0001 5.291 36.36<br>Proto vs. ILA 0.1265 2.491 16.50<br>Proto vs. PC 0.0035 4.897 9.658<br>Fibro vs. ILA 0.9985 0.4237 23.51<br>Fibro vs. PC 0.9997 0.3190 22.37<br>ILA vs. PC >0.9999 0.2138 21.00 | | | | |
| Figure 2N | Brown-Forsythe ANOVA | 3 | 27.81 | 4.88 | 0.0075 |

| | | | | | |
|---|---|---|---|---|---|
| | Welch’s ANOVA | 3 | 9.547 | 5.671 | 0.0168 |
| | **Dunnett's T3 multiple comparisons test**<br><br>**Test** **Adjusted P value** **t** **DF**<br>Proto vs. Fibro 0.0050 3.634 37.24<br>Proto vs. ILA 0.1286 2.483 16.20<br>Proto vs. PC 0.9758 0.6061 2.391<br>Fibro vs. ILA 0.9815 0.6762 21.22<br>Fibro vs. PC 0.1908 2.785 4.313<br>ILA vs. PC 0.1438 2.468 12.99 | | | | |
| Figure 4H | Brown-Forsythe ANOVA | 5 | 69.08 | 46.13 | <0.0001 |
| | Welch’s ANOVA | 5 | 29.85 | 32.32 | <0.0001 |
| | **Dunnett's T3 multiple comparisons test**<br><br>**Test** **Adjusted P value** **t** **DF**<br>CO Proto vs. Ms Proto <0.0001 10.61 32.02<br>CO Fibro vs. Ms Fibro <0.0001 5.504 36.98<br>CO ILA vs. Ms pial 0.0101 3.430 15.52 | | | | |
| Figure 4I | Brown-Forsythe ANOVA | 5 | 37.83 | 17.00 | <0.0001 |
| | Welch’s ANOVA | 5 | 27.29 | 24.86 | <0.0001 |
| | **Dunnett's T3 multiple comparisons test**<br><br>**Test** **Adjusted P value** **t** **DF**<br>CO Proto vs. Ms Proto <0.0001 6.749 36.32<br>CO Fibro vs. Ms Fibro <0.0001 8.619 29.19<br>CO ILA vs. Ms pial 0.0021 4.036 19.00 | | | | |
| Figure 4J | Brown-Forsythe ANOVA | 5 | 64.04 | 22.00 | <0.0001 |
| | Welch’s ANOVA | 5 | 38.42 | 46.78 | <0.0001 |
| | **Dunnett's T3 multiple comparisons test**<br><br>**Test** **Adjusted P value** **t** **DF**<br>CO Proto vs. Ms Proto <0.0001 6.156 31.90<br>CO Fibro vs. Ms Fibro <0.0001 4.844 32.87<br>CO ILA vs. Ms pial <0.0001 7.842 18.01 | | | | |
| Figure 4K | Brown-Forsythe ANOVA | 5 | 63.85 | 26.96 | <0.0001 |

| | | | | | |
|---|---|---|---|---|---|
| | Welch's ANOVA | 5 | 31.05 | 18.21 | <0.0001 |
| | **Dunnett's T3 multiple comparisons test**<br><br>**Test** **Adjusted P value** **t** **DF**<br>CO Proto vs. Ms Proto <0.0001 4.898 35.16<br>CO Fibro vs. Ms Fibro 0.8669 0.6900 33.27<br>CO ILA vs. Ms pial 0.0394 2.750 17.30 | | | | |
| Figure 4L | Brown-Forsythe ANOVA | 5 | 63.03 | 21.98 | <0.0001 |
| | Welch's ANOVA | 5 | 38.44 | 46.63 | <0.0001 |
| | **Dunnett's T3 multiple comparisons test**<br><br>**Test** **Adjusted P value** **t** **DF**<br>CO Proto vs. Ms Proto <0.0001 5.567 31.39<br>CO Fibro vs. Ms Fibro 0.0139 3.010 36.82<br>CO ILA vs. Ms pial 0.0008 4.553 16.83 | | | | |
| Figure 4M | Brown-Forsythe ANOVA | 5 | 59.39 | 20.86 | <0.0001 |
| | Welch's ANOVA | 5 | 26.88 | 16.90 | <0.0001 |
| | **Dunnett's T3 multiple comparisons test**<br><br>**Test** **Adjusted P value** **t** **DF**<br>CO Proto vs. Ms Proto 0.5231 1.244 29.43<br>CO Fibro vs. Ms Fibro <0.0001 8.255 37.89<br>CO ILA vs. Ms pial 0.0068 3.776 13.09 | | | | |
| Figure 4N | Brown-Forsythe ANOVA | 5 | 29.51 | 10.50 | <0.0001 |
| | Welch's ANOVA | 5 | 27.30 | 15.40 | <0.0001 |
| | **Dunnett's T3 multiple comparisons test**<br><br>**Test** **Adjusted P value** **t** **DF**<br>CO Proto vs. Ms Proto 0.0003 4.514 29.79<br>CO Fibro vs. Ms Fibro <0.0001 6.694 30.47<br>CO ILA vs. Ms pial 0.0118 3.294 17.59 | | | | |
| Figure 5E | Brown-Forsythe ANOVA | 2 | 11.49 | 54.05 | <0.0001 |
| | Welch's ANOVA | 2 | 9.748 | 93.92 | <0.0001 |
| | **Dunnett's T3 multiple comparisons test** | | | | |

| | | | | | |
|---|---|---|---|---|---|
| | **Test** | **Adjusted P value** | **t** | **DF** | |
| | Human vs. CO | 0.0380 | 3.257 | 6.763 | |
| | Human vs. Mouse | 0.0016 | 9.699 | 4.138 | |
| | CO vs. Mouse | <0.0001 | 10.61 | 32.02 | |
| Figure 5F | Brown-Forsythe ANOVA | 2 | 6.219 | 15.55 | 0.004 |
| | Welch's ANOVA | 2 | 9.80 | 30.60 | <0.0001 |
| | **Dunnett's T3 multiple comparisons test** | | | | |
| | **Test** | **Adjusted P value** | **t** | **DF** | |
| | Human vs. CO | 0.1362 | 2.500 | 4.734 | |
| | Human vs. Mouse | 0.0230 | 4.732 | 4.109 | |
| | CO vs. Mouse | <0.0001 | 6.749 | 36.32 | |
| Figure 5G | Brown-Forsythe ANOVA | 2 | 6.006 | 34.71 | 0.0005 |
| | Welch's ANOVA | 2 | 9.556 | 42.58 | <0.0001 |
| | **Dunnett's T3 multiple comparisons test** | | | | |
| | **Test** | **Adjusted P value** | **t** | **DF** | |
| | Human vs. CO | 0.0090 | 5.232 | 4.782 | |
| | Human vs. Mouse | 0.0046 | 7.391 | 4.040 | |
| | CO vs. Mouse | <0.0001 | 6.156 | 31.90 | |
| Figure 5H | Brown-Forsythe ANOVA | 2 | 6.556 | 44.90 | 0.0001 |
| | Welch's ANOVA | 2 | 9.779 | 43.14 | <0.0001 |
| | **Dunnett's T3 multiple comparisons test** | | | | |
| | **Test** | **Adjusted P value** | **t** | **DF** | |
| | Human vs. CO | 0.0036 | 6.423 | 4.929 | |
| | Human vs. Mouse | 0.0028 | 8.443 | 4.113 | |
| | CO vs. Mouse | <0.0001 | 4.898 | 35.16 | |
| Figure 5I | Brown-Forsythe ANOVA | 2 | 4.724 | 46.00 | 0.0008 |
| | Welch's ANOVA | 2 | 9.485 | 41.90 | <0.0001 |
| | **Dunnett's T3 multiple comparisons test** | | | | |
| | **Test** | **Adjusted P value** | **t** | **DF** | |
| | Human vs. CO | 0.0073 | 6.516 | 4.293 | |
| | Human vs. Mouse | 0.0039 | 7.706 | 4.012 | |

| | | | | | |
|---|---|---|---|---|---|
|  | CO vs. Mouse <0.0001 5.567 31.39 |  |  |  |  |
| Figure 5J | Brown-Forsythe ANOVA | 2 | 31.93 | 6.732 | 0.0036 |
| | Welch's ANOVA | 2 | 8.10 | 14.55 | 0.0043 |
|  | **Dunnett's T3 multiple comparisons test**<br><br>**Test** **Adjusted P value** **t** **DF**<br>Human vs. CO 0.0047 4.050 11.84<br>Human vs. Mouse 0.0475 2.859 10.39<br>CO vs. Mouse 0.5231 1.244 29.43 |  |  |  |  |
| Figure 5K | Brown-Forsythe ANOVA | 2 | 32.86 | 17.77 | <0.0001 |
| | Welch's ANOVA | 2 | 14.85 | 12.74 | 0.0006 |
|  | **Dunnett's T3 multiple comparisons test**<br><br>**Test** **Adjusted P value** **t** **DF**<br>Human vs. CO 0.0027 4.358 12.49<br>Human vs. Mouse >0.9999 0.02948 10.63<br>CO vs. Mouse 0.0003 4.514 29.79 |  |  |  |  |
